# PUNCH2: explore the strategy for Intrinsically Disordered Protein predictor

**DOI:** 10.1101/2024.10.03.616458

**Authors:** Di Meng, Gianluca Pollastri

## Abstract

Intrinsically disordered proteins (IDPs) lack stable three-dimensional structures, which poses significant challenges for their computational prediction. This study introduces PUNCH2, a novel approach for predicting intrinsically disordered regions (IDRs) in proteins. We address key issues such as the scarcity of comprehensive IDR databases, effective feature extraction, and robust model architecture. By integrating sequences from experimental PDB datasets and fully disordered DisProt sequences for training, PUNCH2 achieves superior prediction accuracy. Various sequence embeddings, including One-Hot, MSA-based, and PLM-based methods, were evaluated, with ProtTrans-based embeddings showing the best performance. The optimal model architecture features multiple convolutional layers, enhancing predictive confidence. PUNCH2 and its lighter variant, PUNCH2-light, demonstrate high efficiency and accuracy in benchmarking against top predictors on the CAID2 dataset, offering promising tools for advancing IDP research and understanding.

## Introduction

Intrinsically disordered proteins (IDPs) are proteins that lack a stable three-dimensional structure and remain highly flexible. This structural flexibility and plasticity enable them to play crucial roles in various biological processes, including signaling, regulation, molecular recognition, and diverse cellular functions. Despite their functional significance, the structure of IDPs is challenging to observe experimentally and predict computationally due to their structural diversity.

It is crucial to note that the training strategy for predictors of intrinsically disordered regions (IDRs) can differ significantly from that of structured regions due to their distinct structural and functional characteristics. In comparison to the prediction of structured regions, which has been a major focus in protein computational studies, IDR prediction, as a relatively new field, faces numerous known and unknown challenges. From a computer science perspective, these challenges can be categorized into the following aspects: a. the availability of high-quality databases; b. effective feature extraction methods and suitable model architectures; c. robust predictor evaluation strategy.

### 1.1 Lack of a Comprehensive IDR Database

Several disordered protein databases exist, such as the general IDR annotation databases DisProt [14][3] which is a community resource annotating protein sequences for intrinsically disordered regions based on literature, and MobiDB [13], which provides annotations from both literature and predictors. Additionally, there are function-specific IDR databases, including DIBS [15], MFIB [8], and PED [9].

While MobiDB and DisProt are widely utilized for analyzing IDRs, they are not typically employed for training IDR predictors. This limitation arises from the inconsistency in the quality of annotations across entire protein sequences. DisProt contains high-quality annotations sourced from literature, but not all residues inside a Disprot sequence are annotated. Conversely, MobiDB offers annotations for entire protein sequences; however, the quality of these annotations varies, containing both experimental results and predictions. These databases can be partially used for predictor evaluation but are not suitable for training predictors. For instance, in the CAID2 challenge [5][6], the dataset defined IDRs based on DisProt annotations and structural regions according to PDB, excluding other regions lacking experimental annotations.

### 1.2 IDR Features Extraction and Model Architecture Design

Protein sequences, composed of amino acids represented by letters, must be converted into numerical representations or matrices for computational analysis. Theoretically, a protein sequence determines its structure and the structure determines its function. In other words, all the information related to protein folding should be encoded in the sequence. Protein sequence embedding techniques transform these sequences into fixed-dimensional matrices with key features encoded. Training models can rely on various features, such as amino acid composition, sequence entropy, hydrophobicity, secondary structure predictions, disorder predictions, predicted structural features (like solvent accessibility), physicochemical properties, and evolutionary information. However, predictors can introduce biases, and experimental data might be unavailable. Therefore, biochemical features and evolutionary information are most commonly used. Biochemical features are static, easily accessible, and interpretable, while evolutionary information, crucial for protein studies, is typically obtained through multiple sequence alignment (MSA), which are based on sets of similar sequences for each query. Nonetheless, there are significant limitations and challenges when using MSA for IDR sequence embedding, primarily concerning the application of MSA results.

The MSA search results yield a list of sequences similar to the query sequences, requiring extra analysis and a more complex model structure to capture hidden information and long-distance dependencies, such as RNN, LSTM and BRNN models. This raises new questions about the optimal representation of MSA results, and consequently the suitable model architectures for learning from these representations. Common combinations include frequency accounting of MSA results, sometimes with one-hot encoding, and the use of SVM, LSTM, or other RNN-based models, as well as hybrid models like the combination of CNN and RNN architectures such as CBRCNN [18]. Another limitation of MSA-based embeddings is their dependency on a large number of similar sequences to form an informative representation. The quality of the representation significantly decreases if there are few or no similar sequences available. This is often the case for highly disordered proteins, as disordered regions are less conserved compared to structural regions. Additionally, MSA-based embedding has limitations due to its computational complexity, time-consuming nature, alignment quality, and ability to capture diverse and context-aware information, making it less suitable for computational protein sequence embedding.

As LLM architectures, particularly Transformer structures [19], have become significant in understanding and representing human-like text from extensive text datasets, there is an emerging trend to apply these models to protein sequences, known as Protein Language Models (PLMs). Compared to MSA, PLMs demonstrate an enhanced capability to capture complex patterns such as sequential information and contextual understanding within extensive datasets. PLMs claim to have acquired knowledge of biochemical, structural, and evolutionary features. Furthermore, PLMs can learn effectively from raw protein sequences or MSA results, producing embeddings in matrix form that are easier to utilize compared to raw MSA results or sequences. PLMs are generally considered faster and more informative than MSA embeddings. However, current evidence does not conclusively demonstrate whether PLMs outperform MSA in IDR prediction, nor identify the optimal embedding method for IDR prediction.

### 1.3 Evaluation of IDR predictors

Evaluating IDR predictors presents several challenges. One major challenge is the potential bias in available datasets, which often favor well-studied proteins and organisms, thereby not fully representing the diversity of protein sequences and structures. The dynamic nature of IDRs further complicates evaluation, as these regions can exhibit context-dependent behavior-being ordered when interacting with partner proteins and disordered when isolated. This variability makes it difficult to assess predictive accuracy consistently. Additionally, choosing performance metrics is crucial but challenging. Common metrics like accuracy, precision, recall, and F1 score might not fully capture the performance of IDR predictors due to the imbalanced nature of datasets, where disordered regions are often less frequent than ordered regions.

## Proposed Solution

The current predictors attempt to employ a predictor-building strategy similar to that used for secondary or tertiary structure prediction but often overlook the distinctions between structured regions and intrinsically disordered regions (IDRs). In light of these challenges, this work aims to concentrate on refining the predictor training process. This includes dataset generation, embedding methods, model architecture, and training strategy, with the objective of exploring and implementing the most suitable approaches to address these challenges and develop a specialized predictor specifically tailored for IDRs.

Due to the lack of fully annotated IDR databases adhering to a single standard, the training set was collected independently. The evaluation/test set s sourced from the CAID2 competition, as it has been used to test all state-of-the-art predictors, allowing for direct comparison with existing models. The most widely used embedding methods are tested, including MSA-based embedding, PLM-based embedding, and their combinations. Additionally, various neural network architectures are examined to determine the optimal combination of neural networks and embedding methods.

As mentioned, MSA search results yield sequences similar to the query sequence. To capture long-distance dependencies and deeply embedded information, model architectures like LSTM, RNN, and CNN-RNN combinations are employed. Common embedding methods include MSA results with one-hot encoding, used with LSTM or other RNN-based models[10], or CNN-RNN combinations like CBRCNN[18][21]. Compared to MSA, PLMs have the ability to capture sequential information and contextual representation within large datasets. Additionally, the embedding results from LLMs are matrices, making it easier for the model to learn patterns compared to raw MSA results. Therefore, theoretically, a simpler model structure may suffice for predictors using LLM embeddings. However, to comprehensively compare and explore the impact of different model structures, the same set of architectures applied to MSA is applied to LLM results.

In this project, only neural network structures are considered, rather than traditional machine learning methods. This decision is based on several factors. First, neural networks such as RNN and LSTM[21][10] have demonstrated superior performance compared to traditional ML methods like SVM[20][22][12]. Second, specific types of neural networks, like CNN, RNN and LSTM, have the ability to capture local and long-distance relationships within data, which is crucial for accurate IDR prediction. Lastly, neural networks generally perform better with large datasets, leveraging their capacity to learn complex patterns and relationships.

The evaluation methods for this project align with those employed in the CAID2[5] competition for two primary reasons. First, using the same test set and evaluation criteria allows for direct comparison with other state-of-the-art IDR predictors. Second, the CAID2 evaluation metrics–AUC, APS, and Maximum F1 score–are particularly effective for assessing IDR predictors. AUC and APS handle imbalanced datasets well, evaluating the model’s ability to distinguish between classes across all thresholds. APS is especially useful for highly imbalanced datasets, where the focus is on the positive class, such as IDR prediction. The F1 score balances precision and recall for a specific threshold, making it valuable for optimizing a general threshold.

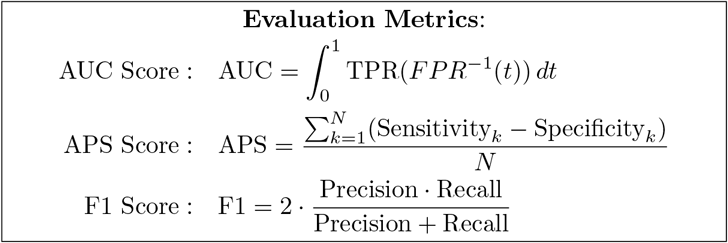

In this project, we aim to build a comprehensive predictor specifically tailored for IDRs by overcoming the current challenges in dataset generation, feature extraction, model architecture, and evaluation. By leveraging state-of-the-art embedding methods and neural network architectures, we develop a robust and accurate IDR predictor that addresses the unique complexities of IDR prediction.

## Experiments

### 3.1 Dataset

To train and evaluate the predictors, we collected several datasets (Table 1). These include *PDB_missing: clstr30, PDB_missing: clstr80*, and *PDB_missing: clstr100*, derived from the PDB [4, 2], and *Disprot_FD*, a fully disordered sequence dataset exported from DisProt [3], for predictor training. Additionally, we used the benchmark evaluation dataset *Disorder_PDB* from the CAID challenge round 2. The detailed dataset collection process and information are shown in Figure 2 and Table 1.

**Table 1.**
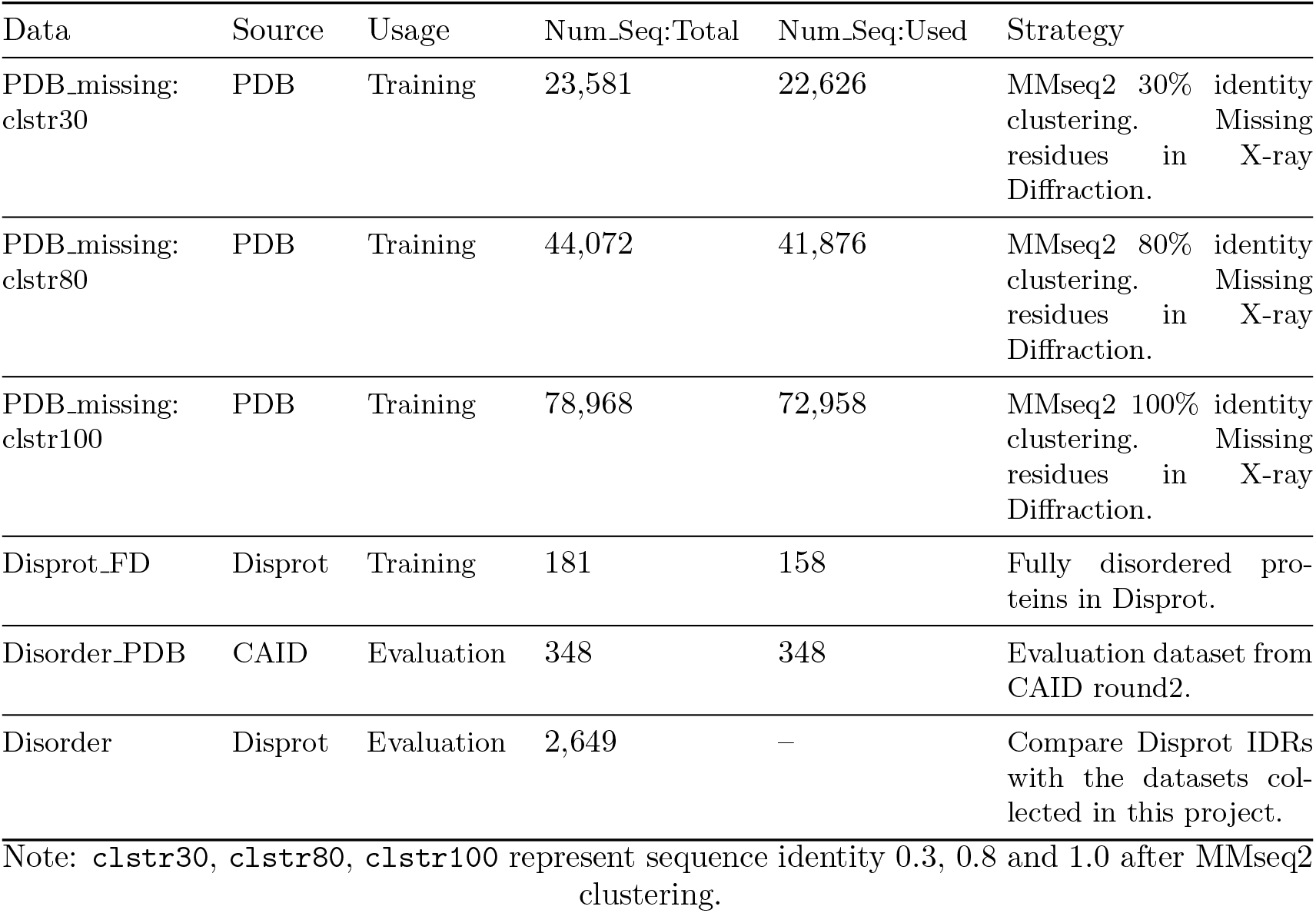
IDR dataset information.

| Data | Source | Usage | Num.Seq:Total | Num.Seq:Used | Strategy |
| --- | --- | --- | --- | --- | --- |
| PDB_missing:<br>clstr30 | PDB | Training | 23,581 | 22,626 | MMseq2 30% identity clustering. Missing residues in X-ray Diffraction. |
| PDB_missing:<br>clstr80 | PDB | Training | 44,072 | 41,876 | MMseq2 80% identity clustering. Missing residues in X-ray Diffraction. |
| PDB_missing:<br>clstr100 | PDB | Training | 78,968 | 72,958 | MMseq2 100% identity clustering. Missing residues in X-ray Diffraction. |
| Disprot_FD | Disprot | Training | 181 | 158 | Fully disordered proteins in Disprot. |
| Disorder_PDB | CAID | Evaluation | 348 | 348 | Evaluation dataset from CAID round2. |
| Disorder | Disprot | Evaluation | 2,649 | – | Compare Disprot IDRs with the datasets collected in this project. |

Note: `clstr30`, `clstr80`, `clstr100` represent sequence identity 0.3, 0.8 and 1.0 after MMseq2 clustering.

#### 3.1.1 IDR_training collection

We collected all PDB entries available as of July 26, 2023, using the query **Structure Determination Methodology = “experimental” AND (Experimental Method = “X-RAY DIFFRACTION” AND Polymer Entity Type = “Protein”)**, resulting in a total of 231,624 entities. Initially, we performed 100% sequence identity clustering to select representative sequences, yielding a dataset of 78,968 PDB sequences. By applying MMseq2[17], these sequences were subsequently clustered into 23,581 groups based on 30% and 44,072 groups based on 80% sequence identity. As shown in Fig 2, this process produced three distinct datasets: *PDB_missing: clstr100*, containing all 78,968 sequences in 23,581 groups; *PDB_missing: clstr80*, comprising 44,072 sequences in 23,581 groups; and *PDB_missing: clstr30*, including only the 23,581 representative sequences from each group. Missing residues identified in X-ray diffraction experiments were annotated as Intrinsically Disordered residues, with no restrictions on minimum or maximum length.

To explain the rationale behind constructing three datasets with different sequence identities: theoretically, a larger dataset with more protein sequences can introduce greater evolutionary diversity, enriching the model training process. This holds true even if sequences within each cluster are similar. However, it remains uncertain to what extent model training benefits from the inclusion of additional similar sequences, or if this inclusion introduces bias towards clusters with more sequences. This concern is particularly relevant post-PLM embedding, given that PLMs are trained on vast amounts of Uniprot sequences. Therefore, *PDB_missing: clstr30* excludes similar sequences (with 30% identity being sufficiently low to assert functional and structural differences between proteins), *PDB_missing: clstr80* contains more similar sequences, and *PDB_missing: clstr100* encompasses the largest number of sequences, exclude only the identical sequences.

Lastly, DisProt only annotates regions of a protein sequence for which its structure is experimentally known. Consequently, some IDR regions may not be correctly annotated due to the lack of experimental evidence. Additionally, *PDB_missing* includes approximately 10% disordered residues within the entire sequences and does not contain any fully disordered sequences. To address this, sequences annotated as 100% disordered in DisProt were extracted, resulting in 181 fully disordered protein sequences (*Disprot_FD*). We compare the features of *Disprot_FD* and *PDB_missing: clstr30* with the ground truth DisProt IDRs. As shown in Figure 1 A & B, DisProt includes a larger percentage of long IDRs compared to *PDB_missing: clstr30*. Both *PDB_missing: clstr30* and *Disprot_FD* exhibit similar amino acid compositions to DisProt IDRs (Figure 1 C & D).

**Figure 1.**
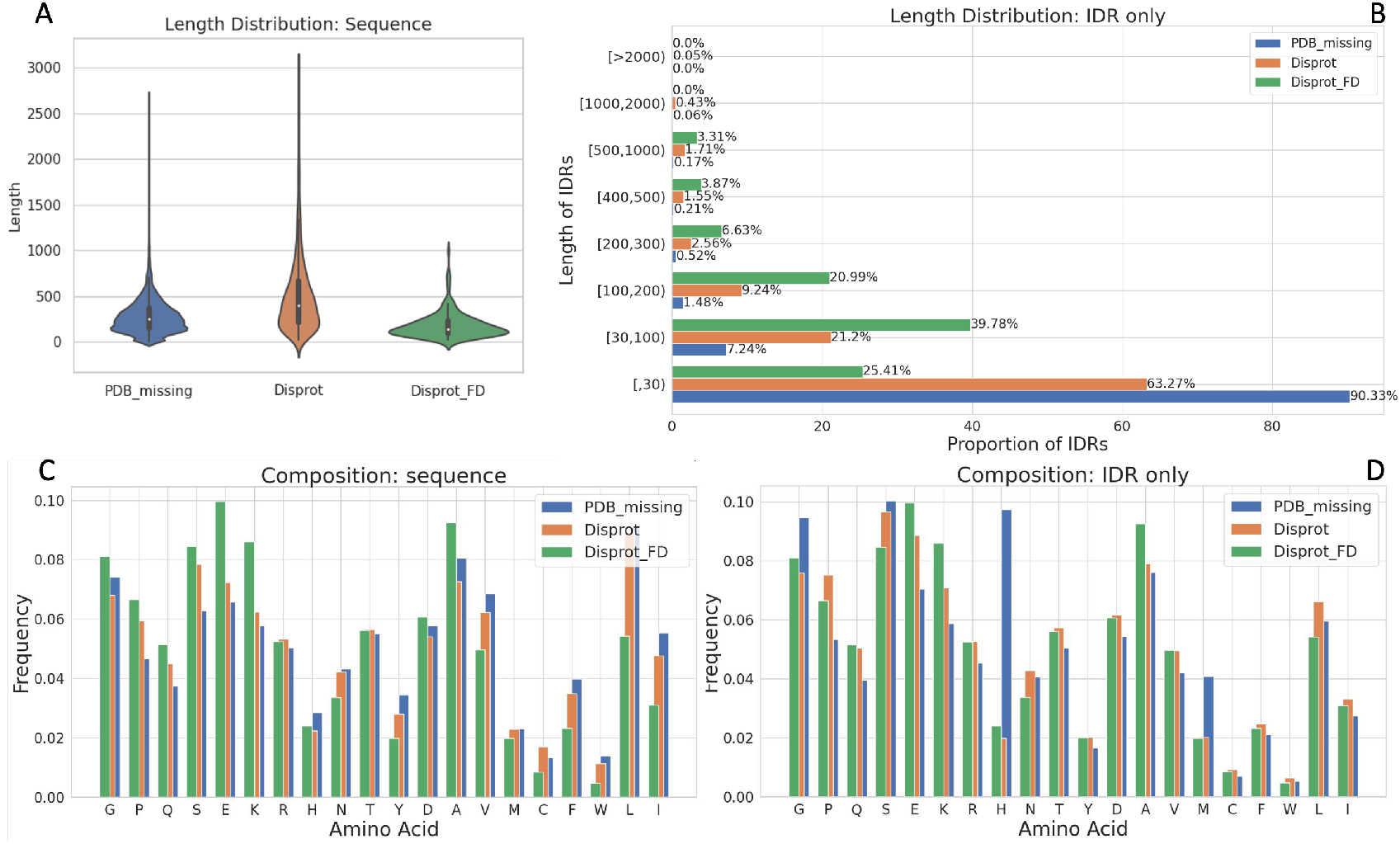
Dataset representation. The graphs present a basic analysis of three datasets: *PDB_missing, Dispot*, and *Disprot_FD*, with *PDB_missing* and *Disprot_FD* serving as our training set. Here, *PDB_missing: clstr30* is used to represent PDB_missing in gerneral. **A**. Violin plots for sequence length distribution: These plots illustrate that *Disprot* has a broader length distribution and a higher percentage of longer sequences, while most *Disprot_FD* sequences are shorter compared to the other two datasets. **B**. Length distribution for IDRs only: For *Disprot_FD*, sequences are considered IDRs since they are fully disordered. The plot reveals that most *Disprot_FD* sequences are longer IDRs, whereas the majority of *PDB_missing* IDRs are shorter, with over 90% of their lengths being less than 30 residues. **C**. Amino acid compositions for sequences: This plot displays the amino acid composition for the sequences in each dataset. **D**. Amino acid compositions for IDRs: This plot shows the amino acid composition for the IDRs in each dataset. The IDR composition plot indicates greater similarity between the three datasets compared to the sequence composition plot.

**Figure 2.**
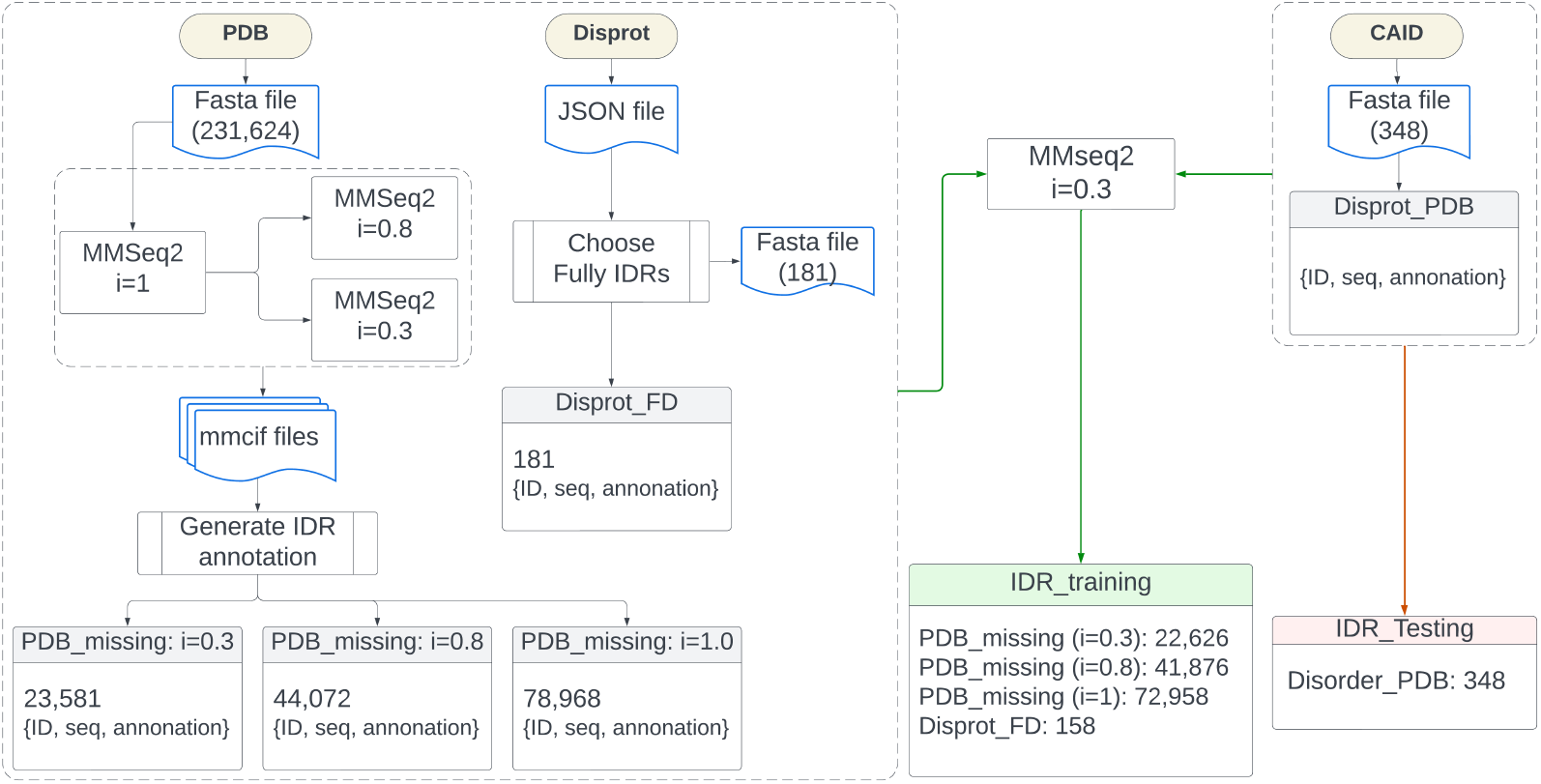
IDR data collection process. In the end, the IDR_Training dataset was searched against the IDR_Training dataset by MMseqs2 with identity=0.3, and exclude the redundant sequences from the IDR_Training.

#### 3.1.2 IDR_testing collection

*Disorder_PDB*, which includes 348 sequences from CAID challenge Round 2 [5], is used as a separate test set. To address the issue of incomplete DisProt annotations, the *Disorder_PDB* dataset was created such that disordered residues are considered positives and only negatives among PDB observed residues are included, leaving other residues unannotated. This dataset is more conservative but is considered more reliable, as it excludes uncertain residues without any structural or disorder annotation.

#### 3.1.3 Final IDR_training & IDR_testing

Finally, to prevent information leakage between the training and test sets, we conducted a search of the training set against the CAID2 *PDB_disorder* dataset using a sequence identity threshold of 30% and excluded all similar sequences from the training set. This precaution ensures that the test set remains independent. As indicated in Table 1, column *Num Seq:Used*, the training set consists of either 72,958, 41,876, or 22,626 sequences from *PDB_missing* clustered into 22,626 groups, supplemented with 158 sequences from *Disprot_FD* representing fully disordered sequences from DisProt. The entire set of 348 sequences from CAID is retained exclusively for our test set.

### 3.2 Sequence Embedding

For sequence embedding, eight embedding methods (Table 2) related to the three most common types of protein sequence embedding methods one-hot encoding, MSA-based embedding, and PLM-based embedding are applied to this project.

**Table 2.** Information for the 8 Embedding methods.

| Name | Input | MAX-length | PLM | Embedding Dim | Description |
| --- | --- | --- | --- | --- | --- |
| One-Hot | Raw sequence | – | – | 21 | feature: 20(common AA) + 1(unusual) |
| MSA-prob | MSA result | – | – | 22 | feature: 20(common AA) + 1(unusual) + 1(Gap) |
| MSA-probAA | MSA result | – | – | 22 | feature: 20(common AA) + 1(unusual) + 1(Gap), set the AA probability in the sequence to 1 |
| MSA-prob-numTemp | MSA result | – | – | 23 | feature: *MSA-prob* + 1 (number of templates from similarity searching) |
| MSA-probAA-numTemp | MSA result | – | – | 23 | feature: *MSA-probAA* + 1 (number of templates from similarity searching) |
| ESM-2 | Raw sequence | 1024 | esm2-t33-650M-UR50D | 1280 | PLM trained on UR90, includes 650M params. |
| MSA Transformer | MSA result | 1024 | esm-msa1b-t12-100M-UR50S | 768 | PLM trained on UR50 + MSA, includes 100M params |
| ProtTrans | Raw sequence | – | ProtT5-XL-UniRef50 | 1024 | PLM trained on UR50, includes 3B params |

Note: The *ESM-2* and *MSA Transformer* models have a sequence length limitation of 1024 tokens, including the start and end tokens (i.e., 1022 sequence tokens plus a start token and an end token).

**One-Hot** encoding represents each amino acid in a protein sequence as a binary vector, with each dimension corresponding to a unique amino acid. This results in a sparse matrix where each amino acid is distinctly represented without any implicit ordinal relationship. Despite its simplicity and limited information content, One-Hot encoding is widely used and often combined with other methods to retain sequence information. In this project, each residue encoded via *One-Hot* encoding will be a 1 *×* 21 vector, with 20 dimensions for common amino acids and 1 for unusual ones.

**MSA**-based embedding leverages the identification of conserved regions among related proteins, capturing essential evolutionary signals and focusing on biologically relevant features like functional domains and motifs. This method enriches the training dataset by searching for similar sequences. Common MSA tools include psiblast[1] and HHblits[16], with HHblits chosen here for its speed. We used HHblits to search against the UniRef30_2020_06 database.

We start with the standard approach of using MSA search results, which counts the frequency of each amino acid at a position (*MSA-prob*), generating a statistical representation of each protein sequence. Additionally, we created three variants: *MSA-probAA*, which sets the probability of the exact residue in the sample sequence to 1; *MSA-prob-numTemp*, which adds a column representing the number of templates from the MSA results to *MSA-prob*; and *MSA-probAA-numTemp*, which adds this column to *MSA-probAA*. As detailed in Table 2, these embeddings have a similar dimension to one-hot encoding but are more densely populated.

**PLMs** utilize deep learning techniques to capture complex patterns and relationships within protein sequences. For PLM-based embedding, we selected *ProtTrans*[7] and *ESM-2* [11], both of which are pre-trained on the UniProt protein sequence database and can extract features from protein sequences. These models produce high-dimensional matrices (1024 and 1280 dimensions, respectively) that represent protein sequences, providing a richer feature set for computational analysis compared to other embedding methods. Additionally, we employed the *MSA-Transformer*, a PLM specifically designed to interpret MSA results, which embeds each amino acid into a 768-dimensional vector. Unlike sequence-based PLMs, the MSA-Transformer utilizes MSA results as inputs, thereby harnessing the evolutionary information present in MSAs.

### 3.3 Model Structure

After selecting the embedding methods, we proceed to determine the model architecture. The choice of model structure is critical, as different embedding methods encode sequence information differently, making certain architectures more suitable for specific embeddings. For instance, some embeddings may benefit from architectures that capture local sequence features, while others may require models capable of identifying long-range dependencies.

Initially, we establish a baseline predictor, **PrepBase**, and a baseline model architecture, **StrucBase. PrepBase** is a static predictor that does not involve any learning process; it is based solely on the amino acid frequency of the IDR region in *PDB_missing: clstr30*. In *PDB_missing: clstr30*, there are 3,972,049 amino acids, with 10% (397,249) located in IDRs. Table 6 and Table S1 (Supplementary) demonstrate that certain amino acids are more prevalent in IDRs than others. Consequently, each specific amino acid is mapped to a corresponding frequency number to generate the prediction result.

The baseline model structure, **StrucBase**, involves learning based solely on the current residue without considering any contextual information. To implement this model structure, we utilize a one-layer convolutional neural network (CNN) with a single channel, a kernel size of one, and *sigmoid* as the activation function, corresponding to *N* = 0 in Figure 3. The primary reason for choosing a CNN is its ability to handle variable-length input sequences.

**Figure 3.**
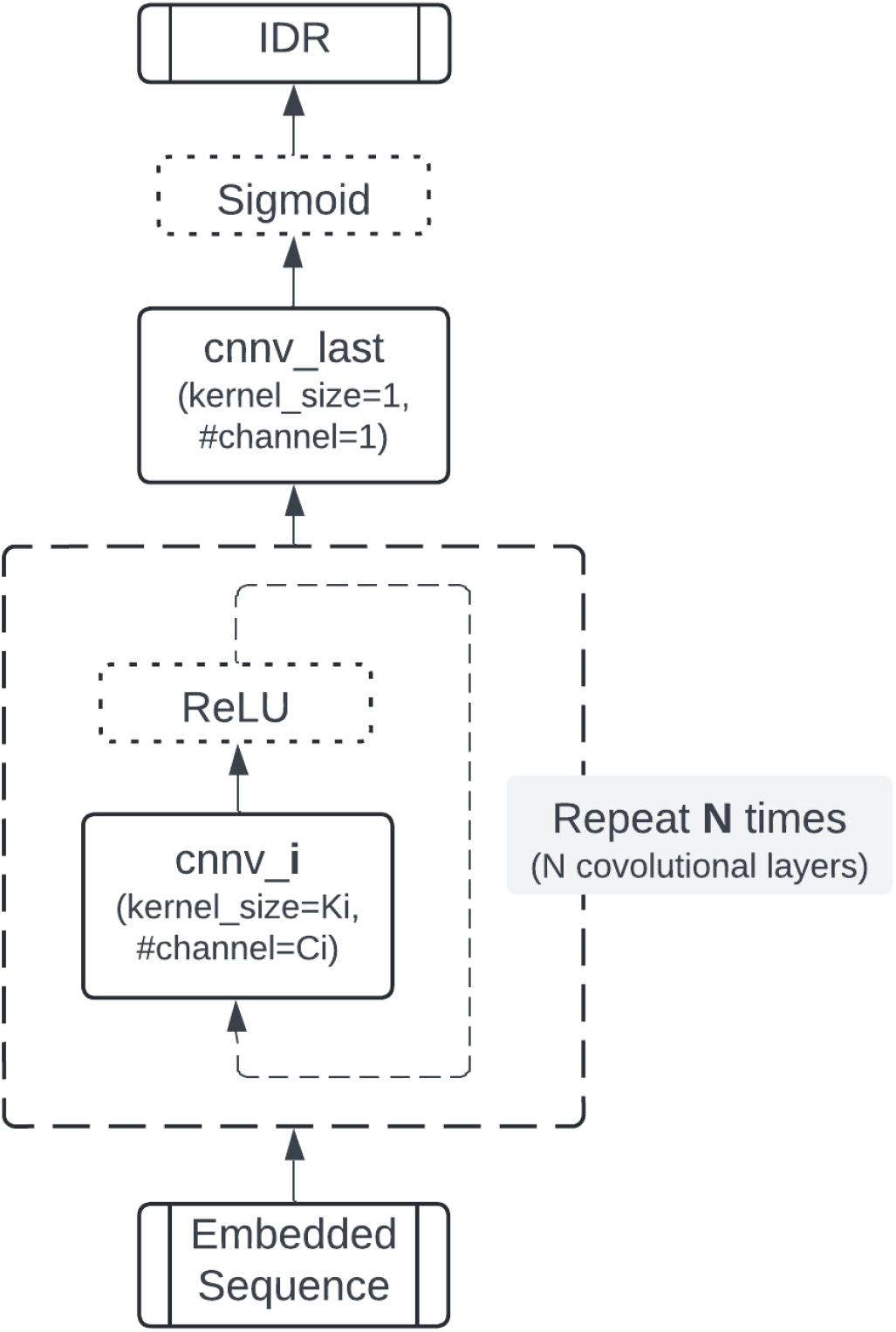
The structure of general CNN-based predictors, where *N* is the total number of Convolutional layers, and *i* is the *i* th Convolutional layer.

Alongside baseline models, we explore RNNs, LSTMs, CNNs, and CBRCNNs for their suitability in IDR and secondary structure prediction. RNNs and LSTMs excel at capturing long-range dependencies in sequential data. CNNs, originally designed for spatial data, are adapted to capture local patterns in sequences through convolutional and pooling layers. Figure 3 outlines our CNN model variations. A modified CBRCNN model, known for its ability to integrate CNNs and RNNs effectively in structured protein prediction tasks, is employed in this study. This architecture combines the local feature extraction capabilities of CNNs with the sequential context understanding of RNNs, making it robust for tasks requiring analysis of sequential data with intricate dependencies. The model variant used here is adapted from the work of Torrisi et al. [18] on protein secondary structure prediction. The structure of our two-stage CBRCNN is depicted in Figure 4, where Stage 1 focuses on predicting IDRs based on embedded sequences, and Stage 2 refines these predictions by incorporating Stage 1’s outputs. Both stages can be trained independently to enhance prediction accuracy. Additionally, Stage 2 of the CBRCNN can be appended to any model architecture. Therefore, it will be incorporated as a secondary stage in our best-performing model architecture.

**Figure 4.**
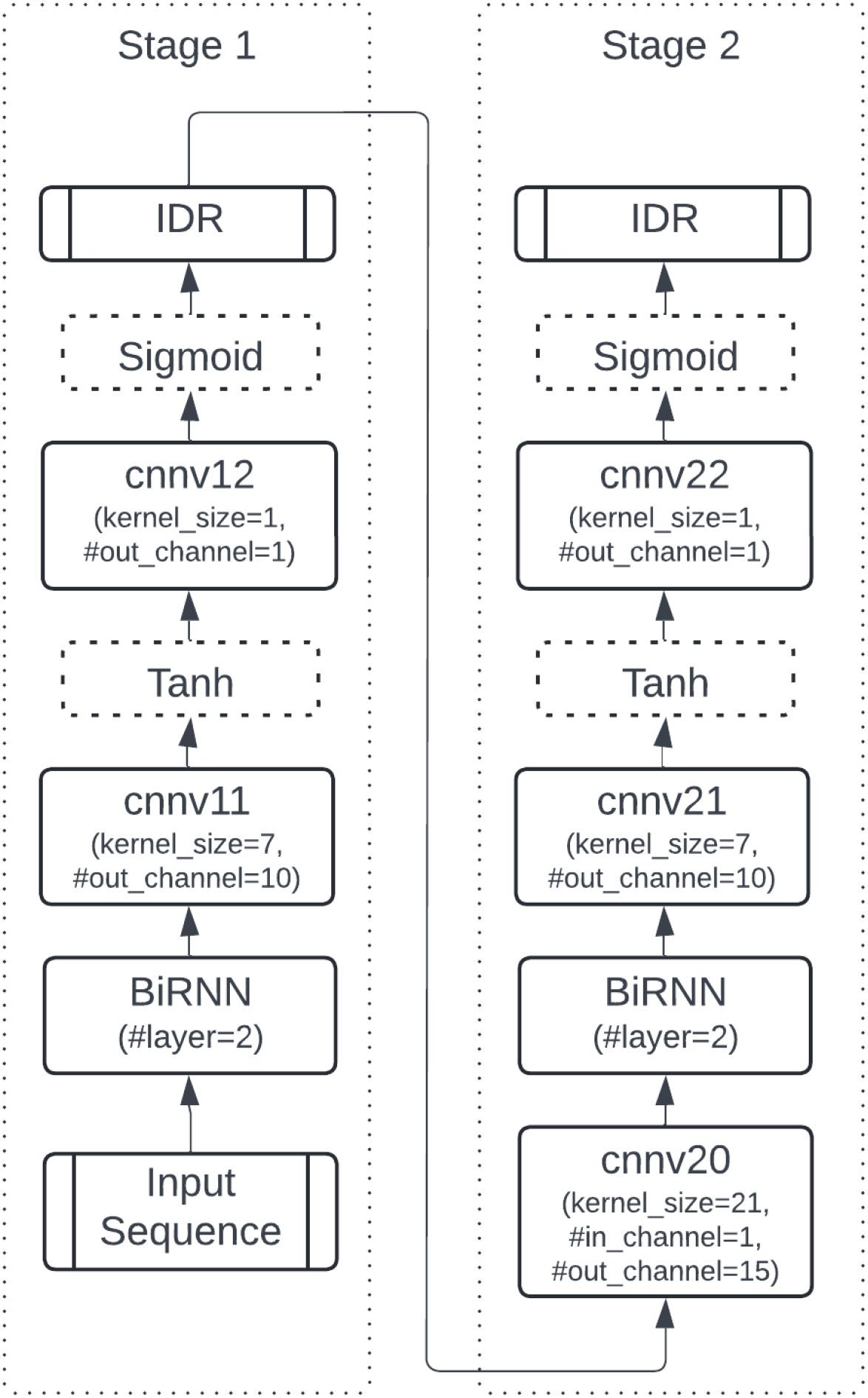
2-stage CBRCNN structure for IDR prediction. The 2 stages can be trained and evaluated separately.

## Training Process & Results

The training process is structured into two phases. `Phase 1` aims to evaluate various embedding methods, model structures, and their combinations. The outcome of Phase 1 consists of several optimal model solutions identified as the most effective combinations of embedding methods and model structures. `Phase 2` implements these optimal configurations on a larger dataset to develop the final predictors, which are subsequently evaluated using the CAID2 test set to ensure their accuracy and robustness.

### 4.1 Phase 1: Embedding methods and Model structures

#### 4.1.1 Performance on Baselines

We begin by employing the baseline model structure **StrucBase** and training on various types of embedded sequences, including One-Hot, MSA-prob, MSA-probAA, MSA-prob-numTemp, MSA-probAA-numTemp, ProtTrans, ESM-2, and MSA Transformer on **StrucBase**. The results are summarized in Table 3. Notably, the AUC score for *One-Hot* encoding is 0.612, slightly outperforming **PredBase**.

**Table 3.**
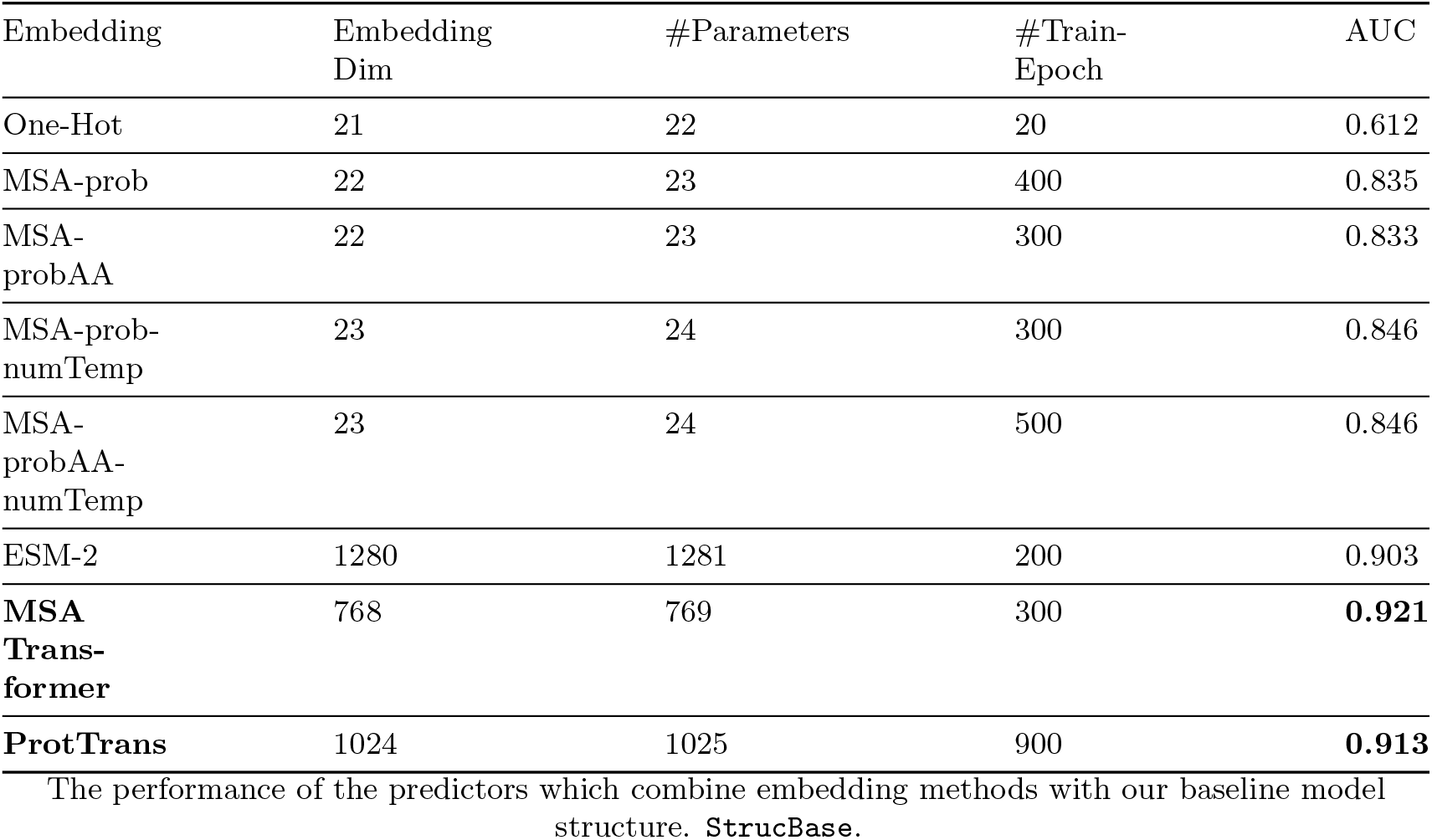
StrucBase: model performance.

Among the MSA-based embedding methods, *MSA-prob* shows no improvement upon altering the amino acid values in the sample sequence (*MSA-probAA*). However, incorporating the number of templates yields a 1.2% improvement in performance.

The PLM-based embedding methods demonstrate superior performance overall. Particularly, *Prot-Trans* and *MSA Transformer* perform exceptionally well, with *MSA Transformer* slightly outperforming *ProtTrans*. It’s important to note that *MSA Transformer* is constrained to sequences shorter than 1022 residues due to its maximum input length of 1024 (including start and end tokens). To handle longer sequences, we applied sequence segmentation with a sliding window approach (*window size* = 1000 residues, *step size* = 500 residues), applying *MSA-Transformer* to each sub-sequence and merging the embeddings while excluding the initial and final 50 residues unless they are true termini. However, the results were suboptimal.

*ProtTrans* and *ESM-2* are similar PLMs trained exclusively on protein sequences, but *ProtTrans* demonstrates better performance than *ESM-2* in IDR embedding. This underscores the effectiveness of PLM-based methods in extracting information from raw protein sequences. With these insights, we proceed to explore more complex model architectures.

#### 4.1.2 Model Architectures

CNN Structure: Shallow vs. Deep We begin with CNNs and explore their hyperparameters using different embedding methods, focusing initially on *One-Hot* encoding, *MSA-prob* embedding, and *ProtTrans* embedding. The hyperparameter tuning process starts with a single CNN layer, then incrementally increases the kernel size, number of channels, and number of layers, evaluating model performance at each step until no further improvement is observed (Figure 3). Once optimal settings are determined, they are applied uniformly across similar embedding methods.

Interestingly, the best-performing hyperparameters for all embedding methods converge to two configurations: **CNN_L11_narrow** and **CNN_L3_wide**. Both models have a similar number of parameters per embedding method (see Supplementary Table S2&S3), with **CNN_L11_narrow** being deeper and narrower, and **CNN_L3_wide** shallower and wider. Despite their structural differences, both configurations yield comparable AUC scores. Notably, in cases such as ProtTrans embedding, as shown in Figure 5, the narrow and deeper CNN structure demonstrates more confidence in its predictions compared to the shallower and wider counterpart, despite similar AUC scores. The detailed structures of **CNN_L11_narrow** and **CNN_L3_wide** are presented in Table 7 and Table 9, respectively.

**Figure 5.**
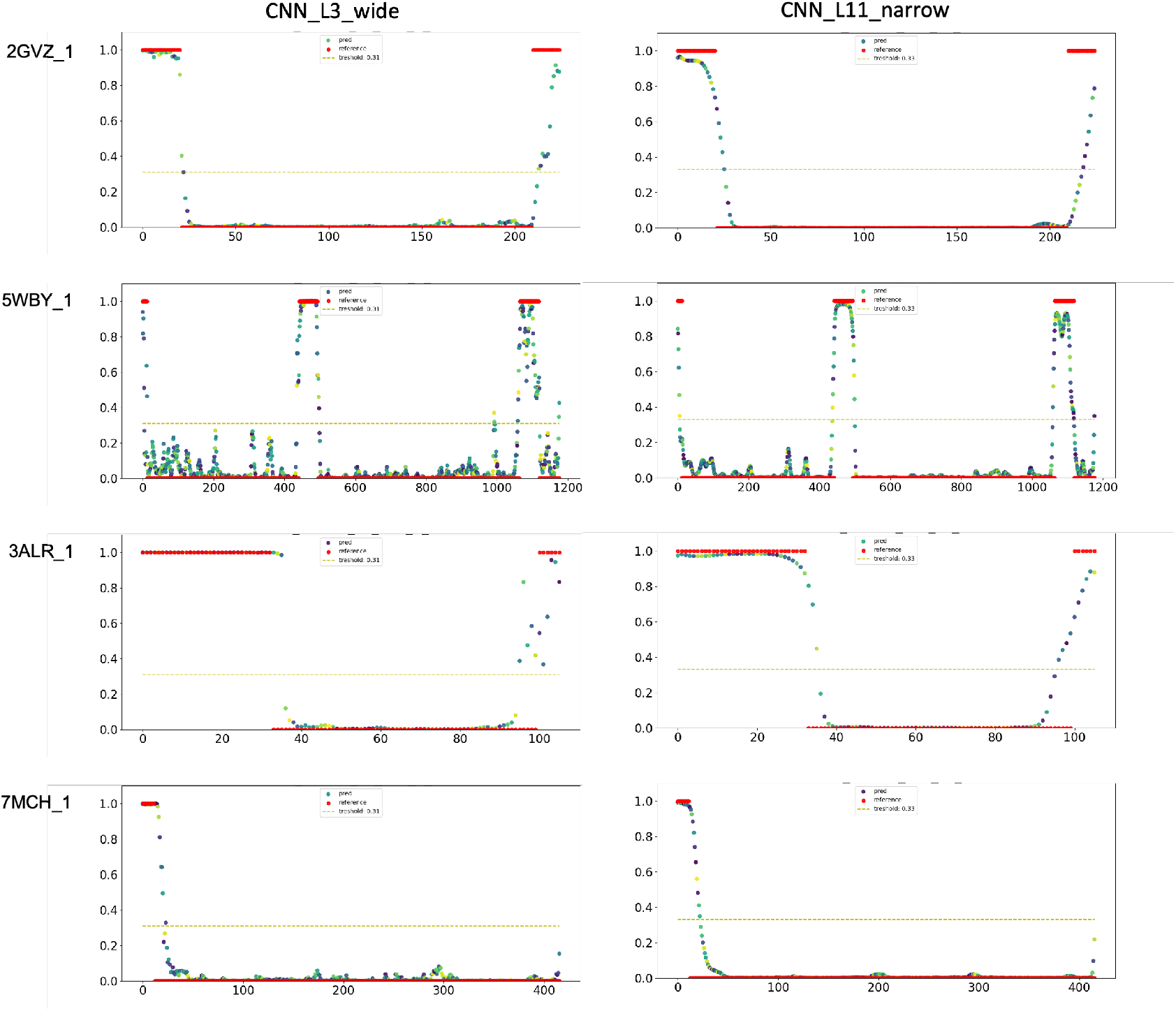
Comparison between shallow (**CNN_L3_wide**) and deeper (**CNN_L11_narrow**) CNN model architectures based on *ProtTrans* sequence embedding, with **CNN_L3_wide** and **CNN_L11_narrow** containing approximately 157K and 180K parameters, respectively. (Red dots: ground truth labels; colourful dots: predictions generated by the corresponding predictor; yellow dish lines: threshold of the predictor.)

RNNs & LSTMs To compare with CNN-based predictors, we applied RNNs and LSTMs using a similar strategy for adjusting hyperparameters across all embedding methods, with particular emphasis on MSA-based embeddings due to their historical performance advantages. However, neither RNNs nor LSTMs matched the performance of **CNN_L11_narrow** across all embedding methods. The best AUC achieved with *ProtTrans* was 0.92 on the test set (with MSA-based embeddings approaching but not reaching 0.9), whereas **CNN_L11_narrow** with *ProtTrans* achieved an AUC of 0.93. Moreover, RNNs and LSTMs required longer training times, making them unsuitable for further training on larger datasets.

CBRCNN CBRCNN, integrating both RNN and CNN components with pre-defined hyperparameters [**?**], yielded slightly better results than RNNs and LSTMs. The best performance using *ProtTrans* reached an AUC of 0.925 in both Stage 1 and Stage 2 (indicating that unlike in secondary structure prediction, the second stage of CBRCNN did not enhance model performance) (Figure 6).

**Figure 6.**
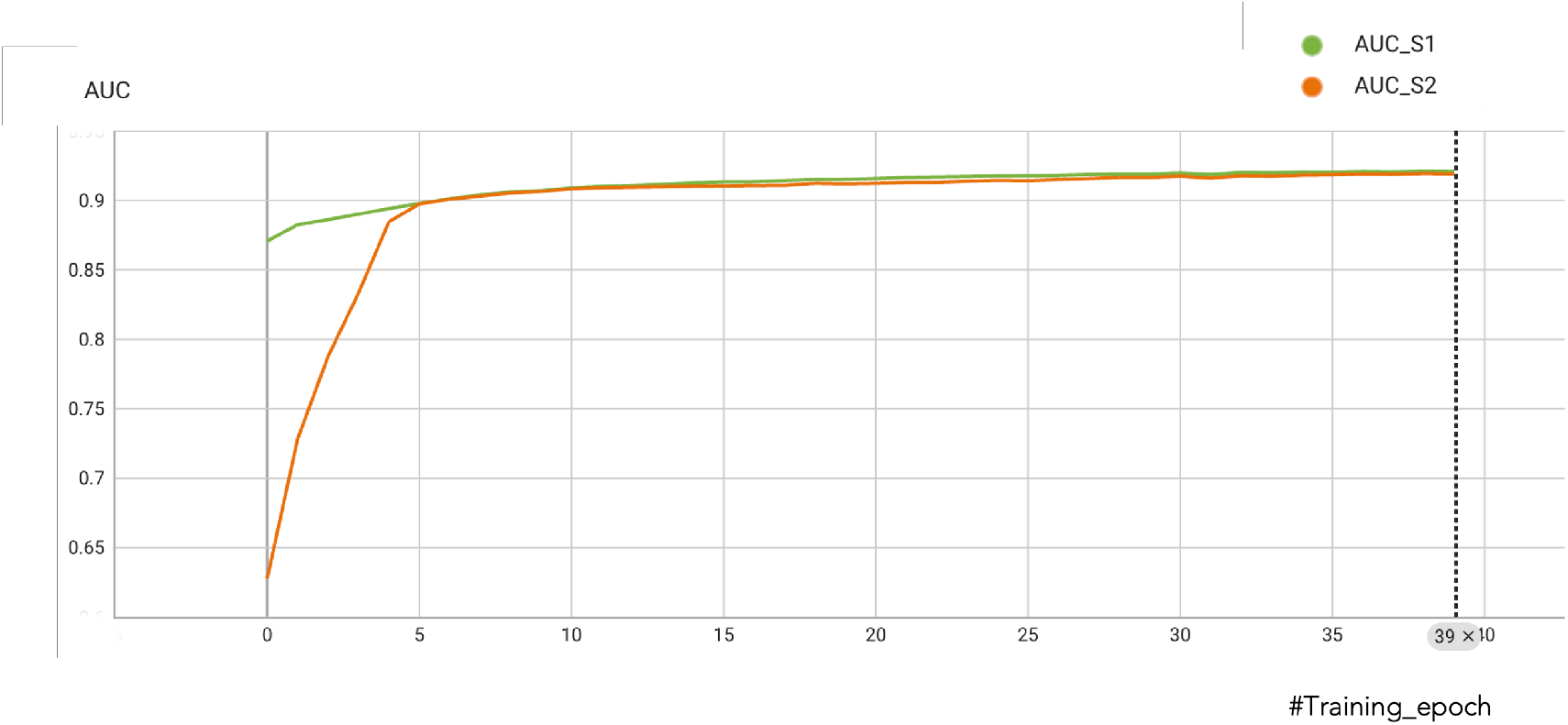
Training progress of CBRCNN, where *AUC_S1* denotes the AUC scores for Stage 1 and *AUC_S2* for Stage 2.

Finally, we augmented **CNN_L11_narrow** with an additional second stage composed exclusively of convolutional layers (Figure 7). However, this modified structure did not enhance the predictor’s performance. Therefore, we have selected **CNN_L11_narrow** for further training.

**Figure 7.**
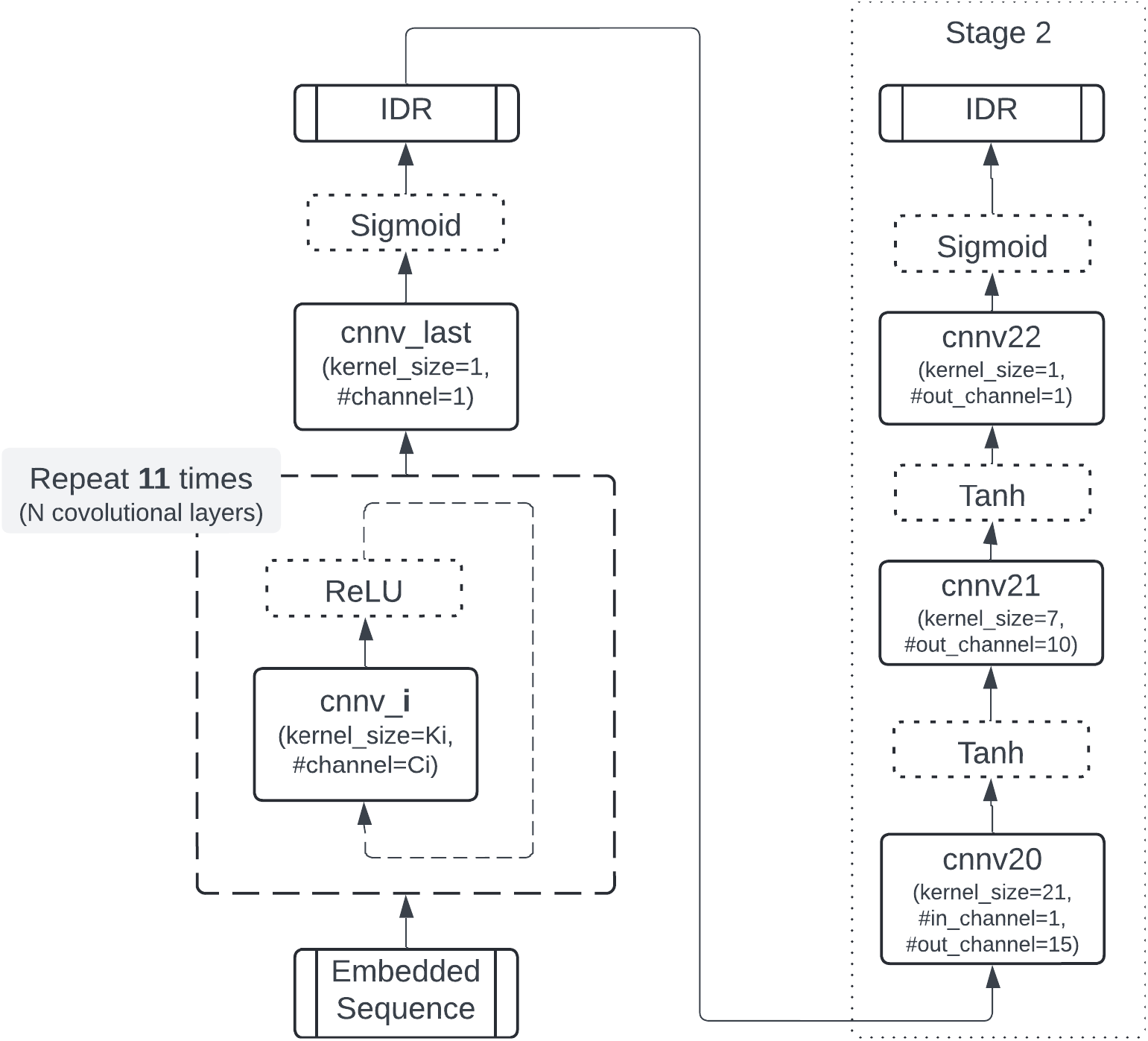
Two-stage CNN structure. Stage 1 represents the best-performing model architecture, **CNN_L3_wide**, while Stage 2 consists of a standalone CNN structure.

### 4.2 Phase 2: Larger dataset and ensemble predictors

In this phase, we evaluate the performance of the **CNN_L11_narrow** model architecture using CAID2 (*Disprot_PDB*) as the test set, while training on datasets excluding sequences similar to *Disprot_PDB*.

#### 4.2.1 Disprot_FD

Initially, we trained models on *PDB_missing: clstr30* alone and on a combined dataset of *PDB_missing: clstr30* and *Disprot_FD*. Table 4 reveals that models trained on *MSA Transformer* and *ProtTrans* show superior performance when incorporating *Disprot_FD*, the predictors’ performance is shown in Table 4 (Predictor 2 vs. 6, 3 vs. 7.). Hence, for subsequent analyses, we utilized the training set comprising *PDB_missing: clstr30* and *Disprot_FD*.

**Table 4.**
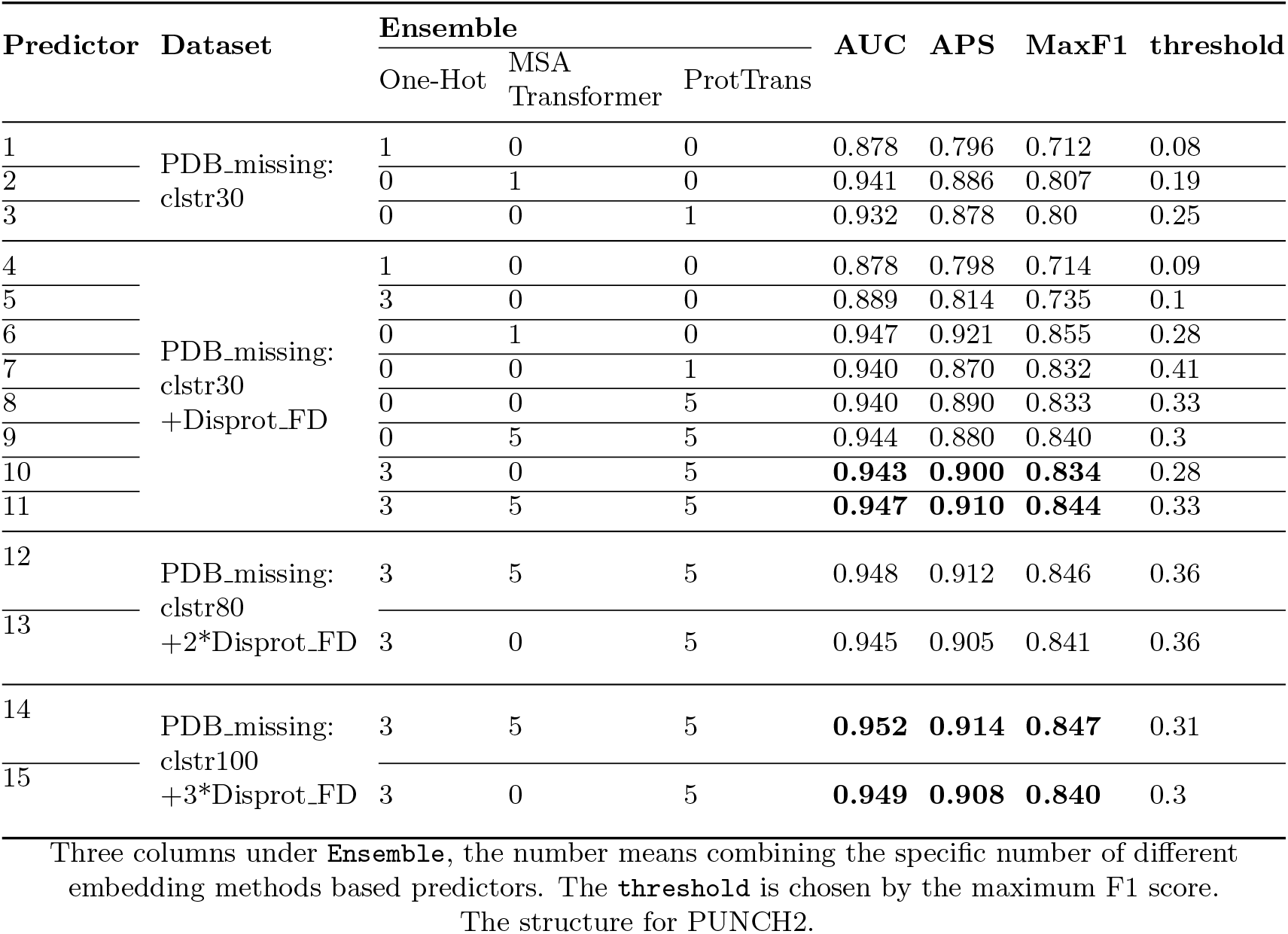
Performance of predictors on Disprot_PDB.

#### 4.2.2 Ensemble predictors

Following individual training on *One-Hot, MSA-Transformer*, and *ProtTrans*, we assessed various combinations of these models. Results indicate that ensemble models consistently outperform individual predictors, with the most effective configuration integrating all three embedding methods.

Subsequently, we conducted 5-fold cross-validation on *MSA-Transformer* and *ProtTrans*, and 3-fold cross-validation on *One-Hot* to construct an ensemble predictor comprising 13 individual predictors. On evaluation against *Disprot_PDB*, this ensemble achieved an AUC of 0.947, APS of 0.91, and maximum F1 score of 0.844 (Table 4). Notably, even without MSA-transformer embedding, the AUC score of the combination of *One-Hot* and *ProtTrans*, Predictor 10, can reach 0.943.

#### 4.2.3 Larger Dataset

To scale our analysis, we transitioned to larger datasets, specifically *PDB_missing:clstr80* and *PDB_missing:clstr100*, and fine-tuned the **CNN_L11_narrow** model architecture accordingly. Our findings indicate the following:

1. Increasing the depth of **CNN_L11_narrow** from 11 to 12 layers (**CNN L12 narrow** Table 8), yielded optimal performance across both larger datasets. The increase in the number of parameters depends on the embedding methods used.
2. To keep a similar level of the importance of fully disordered sequences, we repeat *Disprot_FD* twice for *PDB_missing: clstr80*, and 3 times for *PDB_missing: clstr100*.
3. The ensemble model structure, comprising predictors trained on *PDB_missing:clstr100* augmented with three iterations of *Disprot_FD*, consistently demonstrated the best performance (Table 4).

#### 4.2.4 Final Predictors

We selected two ensemble predictors as our final models. The first, named **PUNCH2**, represents the best-performing ensemble predictor detailed in Table 4, combines 13 single predictors including 3 *One-hot* embedding based predictors, 5 *MSA Transformer* embedding based predictors, and 5 *ProtTrans* embedding based predictors. A drawback of **PUNCH2** is its reliance on five predictors trained on *MSA Transformer* -embedded sequences, which are computationally intensive due to the requirement for multiple sequence alignments (MSA). To mitigate this limitation, we developed a faster variant named **PUNCH2-light**.

**PUNCH2-light** replaces the five *MSA Transformer* -based predictors in **PUNCH2** with five predictors trained on *ProtTrans*-embedded sequences using the **CNN_L3_wide** architecture. Consequently, **PUNCH2-light** consists of three predictors trained on **CNN L12 narrow** with *One-Hot* encoded sequences, five predictors trained on **CNN L12 narrow** with *ProtTrans* embedded sequences, and five predictors trained on **CNN_L3_wide** with *ProtTrans* embedded sequences. Table 5 presents the performance comparison of **PUNCH2** and **PUNCH2-light** with the top 10 predictors from the CAID2 challenge on Disprot_PDB. Both models outperform the other top predictors, demonstrating superior performance metrics, where **PUNCH2** gets AUC, APS, and MaxF1 0.951, 0.914, and 0.847 respectively, and **PUNCH2-light** gets 0.95, 0.912, and 0.845 respectively on CAID2 dataset. Notably, **PUNCH2-light** offers significantly improved computational efficiency compared to **PUNCH2** and other MSA-based predictors.

**Table 5.** Performance comparison of PUNCH2 and Top 10 CAID Predictors on Disprot_PDB.

| Top | Predictor | AUC | APS | MaxF1 |
| --- | --- | --- | --- | --- |
| 1 | SPOT-Disorder2 | 0.949 | 0.928 | 0.86 |
| 2 | AlphaFold-rsa | 0.944 | 0.916 | 0.849 |
| 3 | PredIDR-long | 0.934 | 0.87 | 0.8 |
| 4 | IDP-Fusion | 0.933 | 0.878 | 0.822 |
| 5 | SPOT-Disorder | 0.931 | 0.889 | 0.824 |
| 6 | SETH-0 | 0.93 | 0.893 | 0.83 |
| 7 | AlphaFold-pLDDT | 0.929 | 0.881 | 0.821 |
| 8 | PredIDR-short | 0.927 | 0.859 | 0.79 |
| 9 | metapredict | 0.923 | 0.834 | 0.819 |
| 10 | DeepIDR-2L | 0.922 | 0.858 | 0.794 |
|  | **PUNCH2** | **0.952** | **0.914** | **0.847** |
|  | **PUNCH2-light** | **0.95** | **0.912** | **0.845** |

The numbers from 1 to 10 represent the performance rankings of various predictors on the CAID2 challenge. The last two entries, PUNCH2 and PUNCH2-light, are our newly developed predictors.

## Conclusion

This project aimed to explore the process of building deep learning-based predictors specifically for intrinsically disordered regions (IDRs) and to ultimately develop robust predictors for these regions. We utilized datasets derived from experimental PDB sequences and fully disordered sequences from Disprot (*Disprot_FD*) for training, and CAID2 sequences (*Disprot_PDB*) for benchmarking. The evaluation included various training sets with differing sequence identities (30%, 80%, and 100% from PDB sequences) to determine the optimal configuration for accurate IDR prediction.

Our findings demonstrate that incorporating *Disprot_FD* sequences significantly enhances predictor performance. These fully disordered sequences provide unique features essential for accurately predicting IDRs, in contrast to the structured regions typically represented in PDB sequences. Notably, datasets with 100% sequence identity (*PDB_missing: clstr100*), combined with slightly larger models, outperformed those with lower sequence identities. This suggests that IDRs possess greater sequence diversity than structured regions, thereby enriching the training data, although it may introduce some degree of bias.

Regarding embedding methods, we evaluated One-Hot encoding, MSA-based embedding, and PLM-based embedding techniques. PLM-based embeddings, particularly ProtTrans, consistently outperformed other methods. ProtTrans exhibits stable performance compared to MSA-transformer and slight superiority over ESM2. Combining One-Hot encoding with ProtTrans and optionally MSA-transformer (when available) proved optimal for capturing IDR-specific features. In terms of neural network architecture, deeper models with multiple convolutional layers and approximately 200K parameters achieved the highest AUC scores. Interestingly, deeper networks provided more robust and confident predictions compared to shallower architectures with similar parameters and AUC scores.

Ultimately, we developed two final predictors, PUNCH2 and PUNCH2-light. PUNCH2 utilizes a CNN with 12 layers trained on *PDB_missing: clstr100* incorporating One-Hot, ProtTrans, and optionally MSA-transformer embeddings from *Disprot_PDB*. PUNCH2-light, a faster variant, replaces MSA-transformer embeddings with ProtTrans embeddings in a similar configuration. Both PUNCH2 and PUNCH2-light outperformed top predictors from the CAID2 challenge (Table 5).

*PUNCH2* and *PUNCH2-light* are straightforward and clear-structured predictors, as they solely rely on information extracted from the protein sequence itself. These models do not incorporate additional predictors, secondary structure information, or biochemical features. The rationale for this approach is twofold: firstly, the focus of this work is on exploring the procedure for building an IDR predictor; secondly, theoretically, all necessary information should already be encoded within the sequences. Consequently, for researchers interested in analyzing IDRs, *PUNCH2* and *PUNCH2-light* serve as effective tools for predicting these regions. For those interested in developing IDR predictors, this work provides valuable insights into the detailed process of constructing such models. PUNCH2 and PUNCH2-light are publicly available on GitHub (https://github.com/deemeng/punch2 and https://github.com/deemeng/punch2_light).

## Appendices

### 6.1 PrepBase

Table 6 presents the probability model, `PrepBase`, which specifies the probability distribution for each amino acid. The probabilities are calculated based on the residue composition from the `PDB_missing: clstr30` dataset. This distribution serves as a foundational reference for the model, helping to encode the inherent likelihood of each amino acid’s occurrence.

**Table 6.** Probability model.

| AA | Number | Percentage (%) | AA | Number | Percentage (%) |
| --- | --- | --- | --- | --- | --- |
| A | 30,063 | 0.757 | C | 2,933 | 0.074 |
| D | 22,011 | 0.554 | E | 28,864 | 0.727 |
| F | 8,711 | 0.219 | G | 34,959 | 0.880 |
| H | 32,858 | 0.827 | I | 11,900 | 0.300 |
| K | 23,982 | 0.604 | L | 25,031 | 0.630 |
| M | 14,168 | 0.357 | N | 17,194 | 0.433 |
| P | 21,787 | 0.549 | Q | 15,994 | 0.403 |
| R | 18,600 | 0.468 | S | 39,834 | 1.003 |
| T | 20,605 | 0.519 | V | 17,303 | 0.436 |
| W | 2,294 | 0.058 | Y | 7,041 | 0.177 |
| X | 1,117 | 0.028 |  |  |  |

**PrepBase:** the probability table for each amino acid. The probabilities are generated from the composition of the residues in **PDB\_missing: clstr30**.

### 6.2 Model Hyperparameters

The model structures for three main CNN architectures are detailed below, each with specific configurations and hyperparameters.

#### 6.2.1 CNN_L11_narrow

As shown in Table 7, this model consists of 11 layers with a narrow configuration. The model employs a learning rate of 0.0001. The narrower architecture is designed to focus on extracting detailed features through multiple convolutional layers, allowing the model to capture subtle patterns in the data.

**Table 7.** Model structure of CNN_L11_narrow.

| #layer | #in_channel | #out_channel | kernel_size | stride | padding | activation function |
| --- | --- | --- | --- | --- | --- | --- |
| 1 | #features | 50 | 3 | 1 | 1 | ReLU |
| 2 | 50 | 30 | 3 | 1 | 1 | ReLU |
| 3 | 30 | 30 | 3 | 1 | 1 | ReLU |
| 4 | 30 | 20 | 7 | 1 | 3 | ReLU |
| 5 | 20 | 20 | 7 | 1 | 3 | ReLU |
| 6 | 20 | 20 | 15 | 1 | 7 | ReLU |
| 7 | 20 | 20 | 7 | 1 | 3 | ReLU |
| 8 | 20 | 20 | 7 | 1 | 3 | ReLU |
| 9 | 20 | 10 | 3 | 1 | 1 | ReLU |
| 10 | 10 | 10 | 3 | 1 | 1 | ReLU |
| last | 1 | 1 | 1 | 1 | 0 | Sigmoid |

**learning\_rate** is 0.0001.

#### 6.2.2 CNN L12 narrow

Table 8 outlines the structure of **CNN L12 narrow**, a variant with an additional convolutional layer compared to **CNN_L11_narrow**. This model also uses a learning rate of 0.0001, aiming to further refine feature extraction for a larger dataset through increased depth, potentially capturing more complex hierarchical features.

**Table 8.**
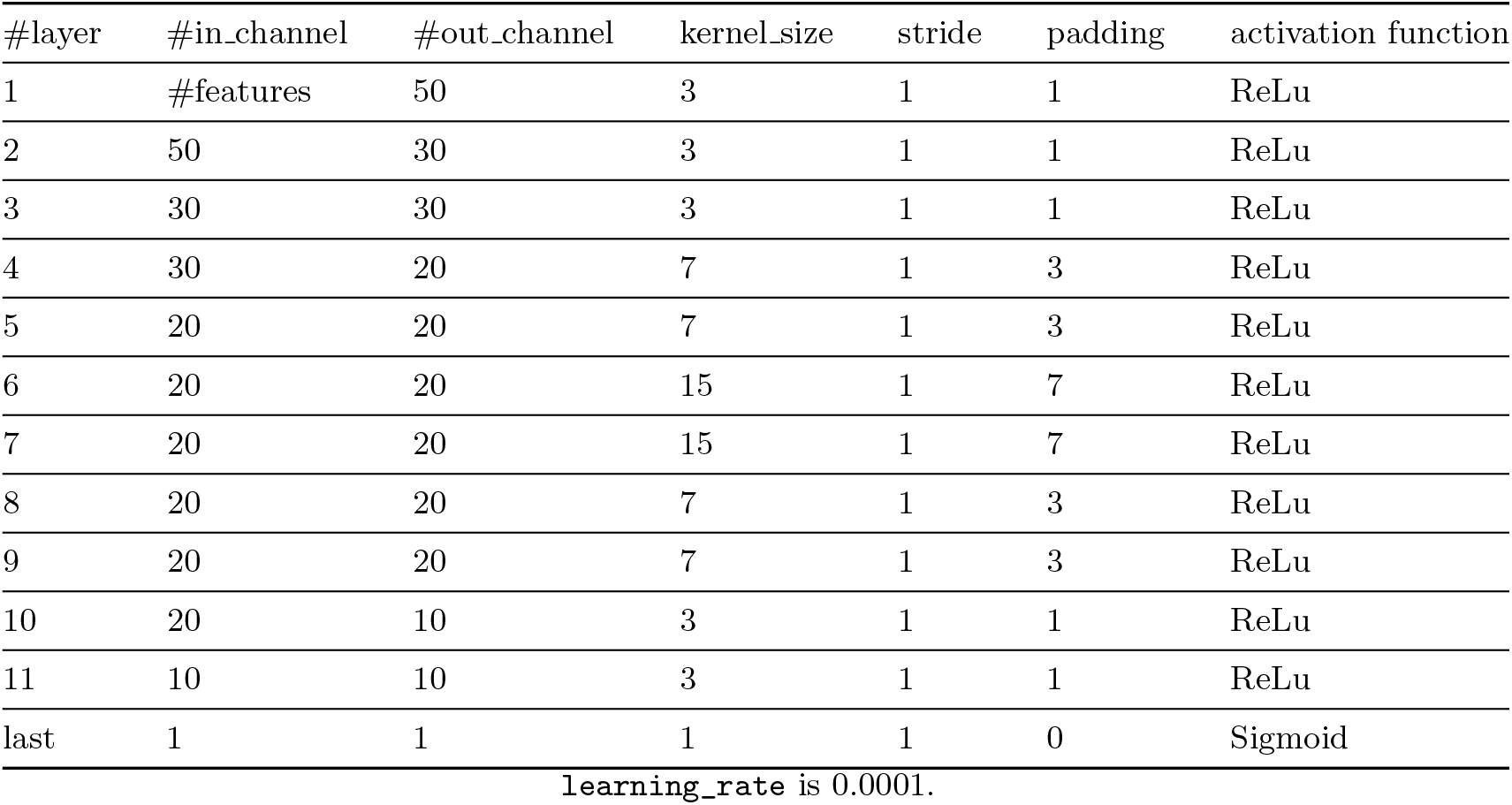
Model structure of CNN L12 narrow.

| #layer | #in_channel | #out_channel | kernel_size | stride | padding | activation function |
| --- | --- | --- | --- | --- | --- | --- |
| 1 | #features | 50 | 3 | 1 | 1 | ReLu |
| 2 | 50 | 30 | 3 | 1 | 1 | ReLu |
| 3 | 30 | 30 | 3 | 1 | 1 | ReLu |
| 4 | 30 | 20 | 7 | 1 | 3 | ReLu |
| 5 | 20 | 20 | 7 | 1 | 3 | ReLu |
| 6 | 20 | 20 | 15 | 1 | 7 | ReLu |
| 7 | 20 | 20 | 15 | 1 | 7 | ReLu |
| 8 | 20 | 20 | 7 | 1 | 3 | ReLu |
| 9 | 20 | 20 | 7 | 1 | 3 | ReLu |
| 10 | 20 | 10 | 3 | 1 | 1 | ReLu |
| 11 | 10 | 10 | 3 | 1 | 1 | ReLu |
| last | 1 | 1 | 1 | 1 | 0 | Sigmoid |

learning\_rate is 0.0001.

#### 6.2.3 CNN_L3_wide

In contrast, Table 9 describes the **CNN_L3_wide** model, which consists of 3 layers but with a wider configuration, meaning each layer has more filters. The wider structure allows for a broader capture of features at each level, facilitating the recognition of diverse patterns within the input data. This model also operates with a learning rate of 0.0001.

**Table 9.** Model structure of CNN_L3_wide.

| #layer | #in_channel | #out_channel | kernel_size | stride | padding | activation function |
| --- | --- | --- | --- | --- | --- | --- |
| 1 | #features | 100 | 3 | 1 | 1 | ReLu |
| 2 | 100 | 50 | 5 | 1 | 2 | ReLu |
| 3 | 50 | 10 | 15 | 1 | 7 | ReLu |
| last | 1 | 1 | 1 | 1 | 0 | Sigmoid |

learning\_rate is 0.0001.

These different configurations highlight the exploration of depth versus breadth in CNN architectures, providing insights into the trade-offs between layer depth and layer width in capturing features from protein sequences.

## Author contributions statement

Must include all authors, identified by initials, for example: D.M. and G.P. conceived the experiment(s), D.M. conducted the experiment(s), D.M. and G.P. analysed the results. D.M. wrote the manuscript and G.P. reviewed the manuscript.

## Acknowledgments

The authors thank Laura Dunne for cooking all the fantastic dinners. This work is supported in part by funds from the Science Foundation Ireland (SFI: # 1636933 and # 1920920).

